# Gut Microbiota and Derived Metabolites Mediate Obstructive Sleep Apnea Induced Atherosclerosis

**DOI:** 10.1101/2024.11.18.624205

**Authors:** Jin Xue, Celeste Allaband, Simone Zuffa, Orit Poulsen, Jason Meadows, Dan Zhou, Pieter C. Dorrestein, Rob Knight, Gabriel G. Haddad

## Abstract

**Background:** Obstructive sleep apnea (OSA) is characterized by intermittent hypoxia/hypercapnia (IHC), affects predominantly obese individuals, and increases atherosclerosis risk. Since we and others have implicated gut microbiota and metabolites in atherogenesis, we dissected their contributions to OSA-induced atherosclerosis.

**Results:** Atherosclerotic lesions were compared between conventionally-reared specific pathogen free (SPF) and germ-free (GF) *ApoE*^-/-^ mice following a high fat high cholesterol diet (HFHC), with and without IHC conditions. The fecal microbiota and metabolome were profiled using 16S rRNA gene amplicon sequencing and untargeted tandem mass spectrometry (LC-MS/MS) respectively. Phenotypic data showed that HFHC significantly increased atherosclerosis as compared to regular chow (RC) in both aorta and pulmonary artery (PA) of SPF mice. IHC exacerbated lesions in addition to HFHC. Differential abundance analysis of gut microbiota identified an enrichment of Akkermansiaceae and a depletion of Muribaculaceae (formerly S24-7) family members in the HFHC-IHC group. LC-MS/MS showed a dysregulation of bile acid profiles with taurocholic acid, taurodeoxycholic acid, and 12-ketodeoxycholic acid enriched in the HFHC-IHC group, long-chain N-acyl amides, and phosphatidylcholines. Interestingly, GF *ApoE*^-/-^ mice markedly reduced atherosclerotic formation relative to SPF *ApoE*^-/-^ mice in the aorta under HFHC/IHC conditions. In contrast, microbial colonization did not show a significant impact on the atherosclerotic progression in PA.

**Conclusions:** In summary, this research demonstrated that (1) IHC acts cooperatively with HFHC to induce atherosclerosis; (2) gut microbiota modulate atherogenesis, induced by HFHC/IHC, in the aorta not in PA; (3) different analytical methods suggest that a specific imbalance between Akkermansiaceae and Muribaculaceae bacterial families mediate OSA-induced atherosclerosis; and (4) derived bile acids, such as deoxycholic acid and lithocholic acid, regulate atherosclerosis in OSA. The knowledge obtained provides novel insights into the potential therapeutic approaches to prevent and treat OSA-induced atherosclerosis.

## Background

Obstructive sleep apnea (OSA) is a respiratory disorder characterized by recurrent upper airway collapse during sleep which leads to inadequate blood gas exchange, namely intermittent hypoxia and hypercapnia (IHC). OSA is a public health concern mainly because of its high prevalence and its wide association with cardiovascular morbidity and mortality [1–5].

OSA is an independent risk factor for atherosclerosis [6–8]. Drager L et al. showed that OSA patients exhibit early signs of atherosclerosis and the level of atherosclerosis correlates with the severity of OSA [7]. Atherosclerosis is a slow, lifelong plaque (i.e. buildup of fats, cholesterol, and other substances) formation and accumulation in the walls of arteries. With plaque buildup, blood vessel walls thicken leading to narrowing of lumens and reduction of blood flow. When these changes are severe enough, such a process can result in life-threatening conditions, such as stroke or myocardial infarction. OSA accelerates atherosclerosis probably by inducing key atherogenic risk factors, such as systemic inflammation, oxidative stress, endothelial dysfunction, elevated systemic blood pressure, platelet activation as well as abnormal glucose and lipid metabolism [6, 8]. Accumulating research evidence suggests that the gut microbiota composition and functionality, via derived metabolites, is also involved in this atherogenic process [9, 10].

The gut microbiota, including bacteria, viruses and fungi, colonizes the gastro-intestinal tract of the host and plays a role in the modulation of immune response, absorption of nutrients, metabolism and development of organ systems in various animals including humans [11, 12]. It is proposed that the gut microbiota may influence the formation of atherosclerosis through different mechanisms: (1) bacterial infection (e.g. lipopolysaccharides and peptidoglycans) can activate the immune system and trigger a harmful inflammatory response that aggravates plaque progression and rupture [13, 14]; (2) cholesterol and lipid metabolism altered by the gut microbiota can modulate the development of atherosclerosis [9]; (3) microbial metabolites can have either beneficial or deleterious effects on atherosclerosis [14]. For example, short-chain fatty acids (SCFAs) [15], bile acids (BAs) [16] and trimethylamine N-oxide (TMAO) [10], have been shown to act as signaling molecules mediating the communications between the gut microbiota and the host to affect the progression of atherosclerosis; and (4) dietary constituents have profound effects on the composition and function of gut microbiota [17–19]. The interplay between diet, microbiota, and microbial metabolites, can impact human health and disease susceptibility, including atherosclerosis [10, 20–23]. Interestingly, we and others have found that the gut microbiota and produced metabolites are altered in different mouse models of OSA [24–26].

However, the mechanistic roles of the gut microbiota and their metabolites in atherosclerosis under a HFHC diet and IHC remain obscure. In the current study, we investigated the contribution of HFHC diet in inducing or promoting atherosclerosis with or without IHC, the changes of gut microbiota and metabolites caused by HFHC diet and IHC, as well as the causal relationship between the gut microbiota and OSA-induced atherosclerosis.

## Methods

### Animals

Ten weeks old male atherosclerosis-prone *ApoE*^-/-^ mice on C57BL/6J background (002052; The Jackson Laboratory, Bar Harbor, ME) [27] were used in the present study. *ApoE* deficiencies were confirmed by polymerase chain reaction (PCR). Germ-free (GF) *ApoE*^-/-^ mice were re-derived in the laboratory of Dr. Sarkis Mazmanian at California Institute of Technology (Caltech) and maintained in the GF core at University of California San Diego (UCSD). GF status was routinely monitored by aerobic and anaerobic cultures, as well as 16S PCR using Zymo Research Quick-DNA Fecal/Soil Microbe Miniprep Kit (D6010, Zymo Research, Irvine, CA). All animal protocols were approved by the Animal Care Committee of the University of California San Diego and followed the Guide for the Care and Use of Laboratory Animals of the National Institutes of Health.

### Diets

Mice were fed either a high fat and high cholesterol diet (HFHC) containing 1.3% cholesterol by weight and 42% fat by Kcal (TD.96121; Envigo-Teklad, Madison, WI) or a regular chow (RC) containing 0.003% cholesterol by weight and 10% fat by Kcal (TD8604; Harlan-Teklad, Madison, WI) for 10 weeks. The body weight of each mouse was measured twice a week. The food intake of the animals in each cage was recorded every week.

### Intermittent Hypoxia and Hypercapnia Treatment

Intermittent hypoxia and hypercapnia (IHC) was administered in a computer-controlled atmosphere chamber (OxyCycler, Reming Bioinstruments, Redfield, NY) as previously described [26]. Mice were exposed to IHC for short periods (∼4 minutes) of 8% O_2_ and 8% CO_2_ separated by alternating periods (∼4 minutes) of normoxia (21% O_2_) and normocapnia (0.5% CO_2_) with 1– 2 minutes ramp intervals, 10 minutes per cycle, 10 hours per day during the light cycle, for 10 weeks. Control mice were under room air (21% O_2_ and 0.5% CO_2_) fed with either the same HFHC or RC diet.

### Quantification of Atherosclerotic Lesions

Atherosclerosis was quantified by computer-assisted image analysis (ImageJ, NIH Image) [28] in Sudan IV-stained en face preparations of the aorta and pulmonary arteries (PA) as previously described [26]. The mouse hearts were perfused with 4% paraformaldehyde. The entire aorta, pulmonary root, and left and right PAs were dissected out and stained with Sudan IV. The magnitude of lesion was presented by the percentage of Sudan IV-stained area to the total area of the blood vessel examined. Images of the aortic arch were cropped from the rest of the aorta (aortic trunk) by measuring the same distance from the bifurcation to the trunk using photo-editing software (Adobe Photoshop CS6, Adobe Systems Inc., San Jose, CA). All the measurements were done by blinded investigators.

### Microbiome Analysis

Fecal samples were collected consistently between 9AM and 11AM (ZT3-ZT5) on collection days and immediately stored at −80 °C until the end of the study. We chose to collect samples at ZT3-ZT5 due to a concomitant circadian study from our group indicating it was the time of greatest microbiome composition differences between IHC and Air [29, 30]. Samples were then prepared for sequencing and analysis following the Earth Microbiome Project standard protocols (http://www.earthmicrobiome.org/protocols-and-standards/16s) [31]. The V4 region of 16S rRNA gene was sequenced using the primer pair 515f to 806r with Golay error-correcting barcodes on the reverse primer. After processing, raw sequence data was uploaded to Qiita (QIITA #11548) [32] and processed using the Deblur workflow (v2021.09) [33] with default parameters into a BIOM format table with amplicon sequencing variants (ASVs). The BIOM table was processed through QIIME 2 (version 2024.2) [34]. Greengenes2 was used for phylogeny and taxonomy [35]. Datasets were rarified to 10,200 reads to control for sequencing effort. Robust principal component analysis (RPCA) [36] and Weighted UniFrac [37] beta diversity distances were used to evaluate and compare microbiome compositional differences. Significance was tested using PERMANOVA [38] and linear mixed effects models [39] (generic equation: variable of interest ∼ host age * diet-exposure group; random effect = host_subject_id). Differential abundance analysis on the feature table collapsed to family level was performed using ANCOM-BC2 [40, 41] and RPCA ranks and visualized using Qurro [42]. Time-informed dimensionality reduction for longitudinal microbiome studies (TEMPoral TEnsor Decomposition = TEMPTED) was also used to identify ASV of interest [43]. TEMPTED dimensionality reduction flattens all information from all timepoints for each mouse to a single point for comparison. Data was visualized using EMPeror [44] and custom python scripts. Log ratios are used instead of relative abundances due to their increased replicability across studies [45, 46].

### Untargeted Metabolomics Analysis

Untargeted liquid chromatography coupled with tandem mass spectrometry (LC-MS/MS) was used to examine the fecal samples as previously described [25]. Briefly, the samples were analyzed on a Vanquish ultrahigh-performance liquid chromatography (UHPLC) system coupled to a Q Exactive mass spectrometer (Thermo Fisher Scientific, Bremen, Germany). Chromatographic separation was achieved with a C18 core shell column (Kinetex, 50 by 2 mm, 1.7-µm particle size, 100-Å pore size; Phenomenex, Torrance, CA). Acquired raw spectra were converted to the open format mzML using MSConvert [47]. Feature detection and extraction was performed via MZmine 4.2 [48]. Briefly, mass detection was performed with MS1 and MS2 noise levels set to 5E4 and 1E3 respectively. Chromatogram builder parameters were set at 5 minimum consecutive scans, 1E5 minimum absolute height, and 10 ppm for *m/z* tolerance. Smoothing was applied before local minimum resolver, which had the following parameters: chromatographic threshold 85%, minimum search range RT 0.2 min, minimum ratio of peak top/edge 1.7. Then, 13C isotope filter and isotope finder were applied. Features were aligned using join aligner with weight for m/z set to 80 and retention time tolerance set to 0.2 min. Features not detected in at least 2 samples were removed before performing peak finder. MetaCorrelate and ion identity networking were performed before exporting the final feature table. The GNPS export function was used to generate the final feature table containing the peak areas and the .mgf file necessary for downstream analysis. Feature based molecular networking (FBMN) [49] was performed in GNPS2 (https://gnps2.org/status?task=0ec6adc1ae534ba3bfcceb9283e3a920).

### Omics Integration

Joint-RPCA is uniquely suited to understanding sparse and compositional data as seen in the final time point of both microbiome and metabolome data. This unsupervised machine learning method for multi-omics data applies robust-centered-log-ratio transformation (rclr), to center the data around zero and approximate a normal distribution, to each matrix [50]. Rclr handles the sparsity without requiring imputation. The joint factorization used builds on the OptSpace matrix completion algorithm, which is a singular value decomposition optimized on a local manifold. For each matrix, the observed values were only computed on the non-zero entries and then averaged, such that the shared estimated matrices were minimized by gradient descent and then optimized across all matrices. To ensure consistency of estimated matrices rotation, the estimated shared matrix and the matrix of shared eigenvalues across all input matrices were recalculated at each iteration. To prevent overfitting, cross-validation of the reconstruction was performed. Samples were randomly assigned to training (80%) or test (20%) sets. Minimization was performed on only the training set data. The test set data were then projected into the same space using the training set data estimated matrices and the reconstruction of the test data was calculated. The correlations of all features across all input matrices were then calculated from the final estimated matrices.

### Statistical Analysis

Lesion data were presented as means ± standard error of the mean (SEM). One-way ANOVA followed by Tukey’s multiple comparison test or Student’s t-test was employed and *p* < 0.05 was considered statistically significant. Metabolomics data were imported in R 4.2.2 (The R Foundation for Statistical Computing, Vienna, Austria) for downstream data analysis. Briefly, quality control samples were used to check data quality and blank subtraction was used to remove noisy features. Features with near zero variance were removed using the package ‘caret v 6.0’. Multivariate analysis was conducted using the package ‘mixOmics v 6.22’ [51]. Principal component analysis (PCA) and partial least square discriminant analysis (PLS-DA) were performed on the peak areas after robust center log ratio transformation (rclr) via the package ‘vegan v 2.6’. For the unsupervised PCA models, PERMANOVA was used to evaluate group centroid separation while to evaluate the performance of the PLS-DA models a 5-folds cross-validation was used. Variable importance (VIP) scores were calculated per feature and features with VIPs > 1 were considered as significant. ANOVA followed by Tukey’s HSD test, for FDR correction, was used to investigate group differences.

## Results

### IHC exacerbates atherosclerotic plaque formation under the HFHC diet

To investigate the impact of high fat and high cholesterol (HFHC) and IHC on atherosclerotic lesion development, SPF *ApoE*^-/-^ mice were fed with either a HFHC diet or regular chow (RC) diet for 10 weeks. Animals were also exposed to either room air (Air) or intermittent hypoxia/ hypercapnia (IHC) to study the potential atherogenic effect of OSA. HFHC-treated mice exhibited significantly more atherosclerotic lesions than those fed with RC in room air (Figure 1). Specifically, the aorta (HFHC-Air 8.1±0.76% vs RC-Air 1.0±0.27%, p<0.001), the aortic arch (HFHC-Air 16.6±2.00% vs RC-Air 2.3±0.55%, p<0.001), and the pulmonary artery (HFHC-Air 12.2±1.51% vs RC-Air 0.2±0.06%, p<0.001) were all affected. Ten-week IHC exposure further aggravated atherogenesis in the aorta, aortic arch, and PA, as compared to Air controls in the presence of HFHC (Aorta, HFHC-IHC 13.8±0.96% vs HFHC-Air 8.1±0.76%, p<0.001; Aortic arch, HFHC-IHC 28.5±1.88% vs HFHC-Air 16.6±2.00%, p<0.001; PA, HFHC-IHC 28.9±2.81% vs HFHC-Air 12.2±1.51%, p<0.001)(Figure 1), indicating that IHC works synergistically with HFHC to influence atherosclerosis.

**Figure 1.**
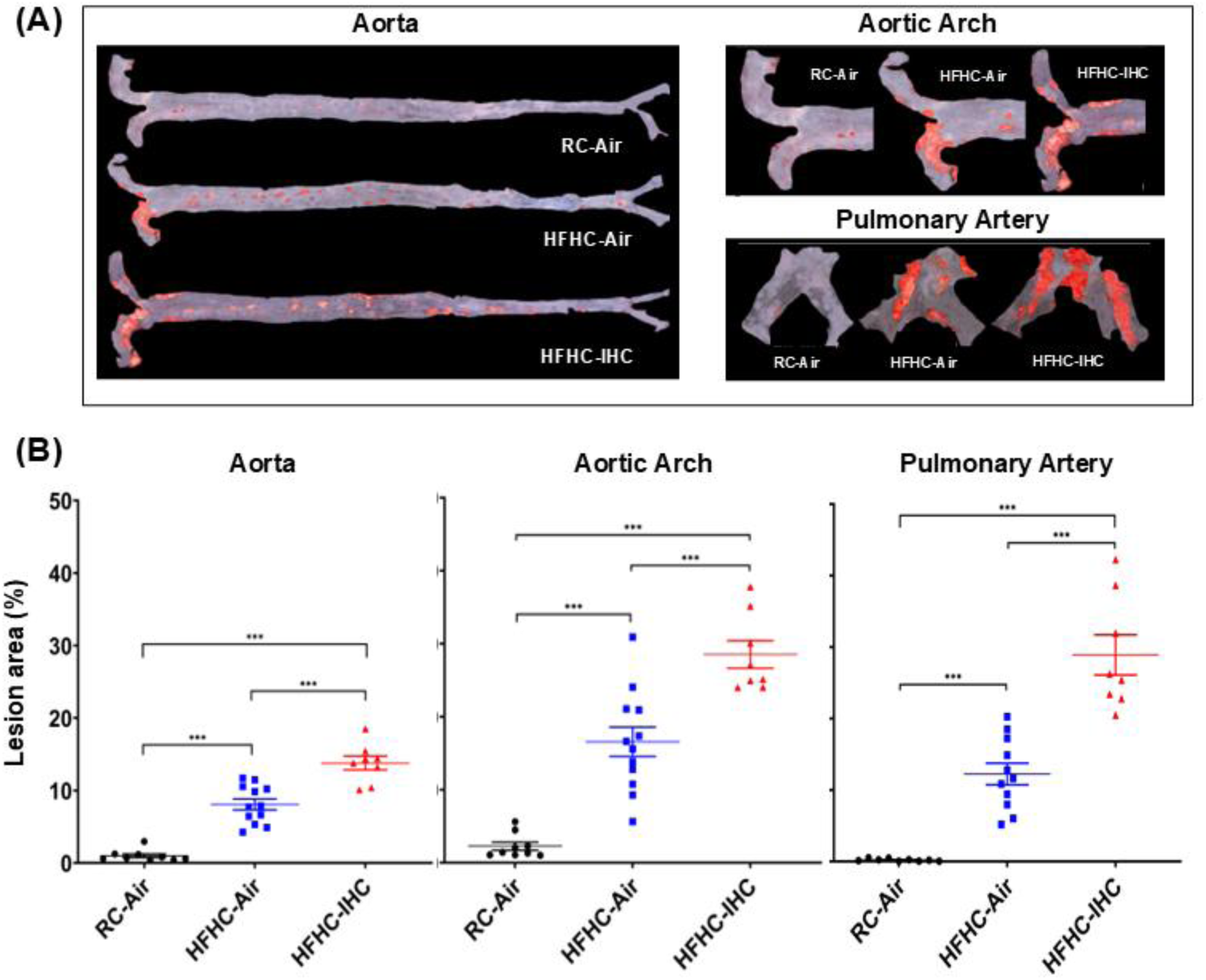
Atherosclerotic lesions after 10-week of HFHC diet with or without IHC in SPF *ApoE*^-/-^ mice. (A) Representative Sudan IV-stained images of lesions. (B) The en-face lesions were quantified as the percentage of lesion area in the total area of the blood vessel examined. HFHC diet increased lesion formation and IHC further accelerated atherosclerotic progression in the presence of HFHC. Data are presented as means ± SEM. Statistical significance tested via One-way ANOVA followed by Tukey’s multiple comparison test. Significance: *** p < 0.001.

To evaluate lesion development with diet and environmental exposure, we measured body weight and food consumption. Not surprisingly, mice on the HFHC diet had significantly higher weight gain than those in the RC group (Supplementary Figure S1A). Intriguingly, IHC exposure decreased body weight gain compared to their Air controls (Supplementary Figure S1A), along with increased lesion formation. Of note, *ApoE*^-/-^ mice consumed more RC food than HFHC food throughout the whole treatment period. IHC stress reduced food intake relative to Air controls under the same HFHC diet condition, starting the week 3 of the treatment (Supplementary Figure S1B).

### Both HFHC and IHC alter the gut microbiota composition

To find out the changes of gut microbiota composition by HFHC and IHC, serial fecal samples from all mice were collected every 3-4 days during the study (10 weeks) and analyzed by 16S V4 amplicon sequencing.

A beta diversity metric based on robust center log ratio (rclr) transformed data, RPCA, was used to examine gut microbial composition (Figure 2A). While groups were not significantly different at the initial time point when all animals were on RC, differences accumulated over time. Strong effects of both diet and experimental exposure type were observed (RC-Air vs HFHC-Air, PERMANOVA p < 0.001, pseudo-F statistic = 295; RC-Air vs HFHC-IHC, PERMANOVA p < 0.001, pseudo-F statistic = 380; HFHC-Air vs HFHC-IHC, PERMANOVA p < 0.001, pseudo-F statistic = 22). Other traditional beta diversity metrics, such as weighted UniFrac, supported these observations (Supplementary Figure S2A). We also saw significant and clear separation between groups when using the TEMPoral TEnsor Decomposition (TEMPTED) beta diversity metric, which captures within-subject correlation and temporal structures in longitudinal datasets better than other methods [43] (Figure 2B). Additionally, examination of alpha diversity metrics also revealed strong effects of diet on microbiome composition, especially for observed features and Faith’s PD, which measures biodiversity (Supplementary Figure S2B-D). Differentially abundant bacterial families identified by ANCOM-BC2 were shown in Figure 2C (IHC vs Air, all on HFHC) and Figure 2D (HFHC vs RC, all in room air). Key overlapping families with opposing enrichments are Muribaculaceae and Akkermansiaceae (Figure 2C-D, bold and star). Based on our previous work [25, 52] as well as differential abundance results from this dataset using RPCA rankings (Figure 2A), TEMPTED rankings (Figure 2B), and ANCOM-BC discriminant features (Figure 2C-D), we examined a relevant natural log ratio comparing reads from ASV belonging to Muribaculaceae (formerly known as S24-7) and Akkermansiaceae families (Figure 2E). This log ratio revealed significant differences among all three groups over time when using linear mixed effects models. Mice on HFHC diet and exposed to IHC conditions were the group most likely to have more Akkermansiaceae than Muribaculaceae, especially by the end of the experiment (Figure 2F). In addition, a log ratio based on the top and bottom TEMPTED differentially ranked ASV for Axis 1 was created, which separated IHC from Air condition, regardless of diet (Supplementary Figure S2E and Supplementary Table S1). A log ratio based on the top and bottom TEMPTED differentially ranked ASV for Axis 2 was also created, which did a better job of separating regular chow from HFHC diet, regardless of exposure type (Supplementary Figure S2F and Supplementary Table S2).

**Figure 2.**
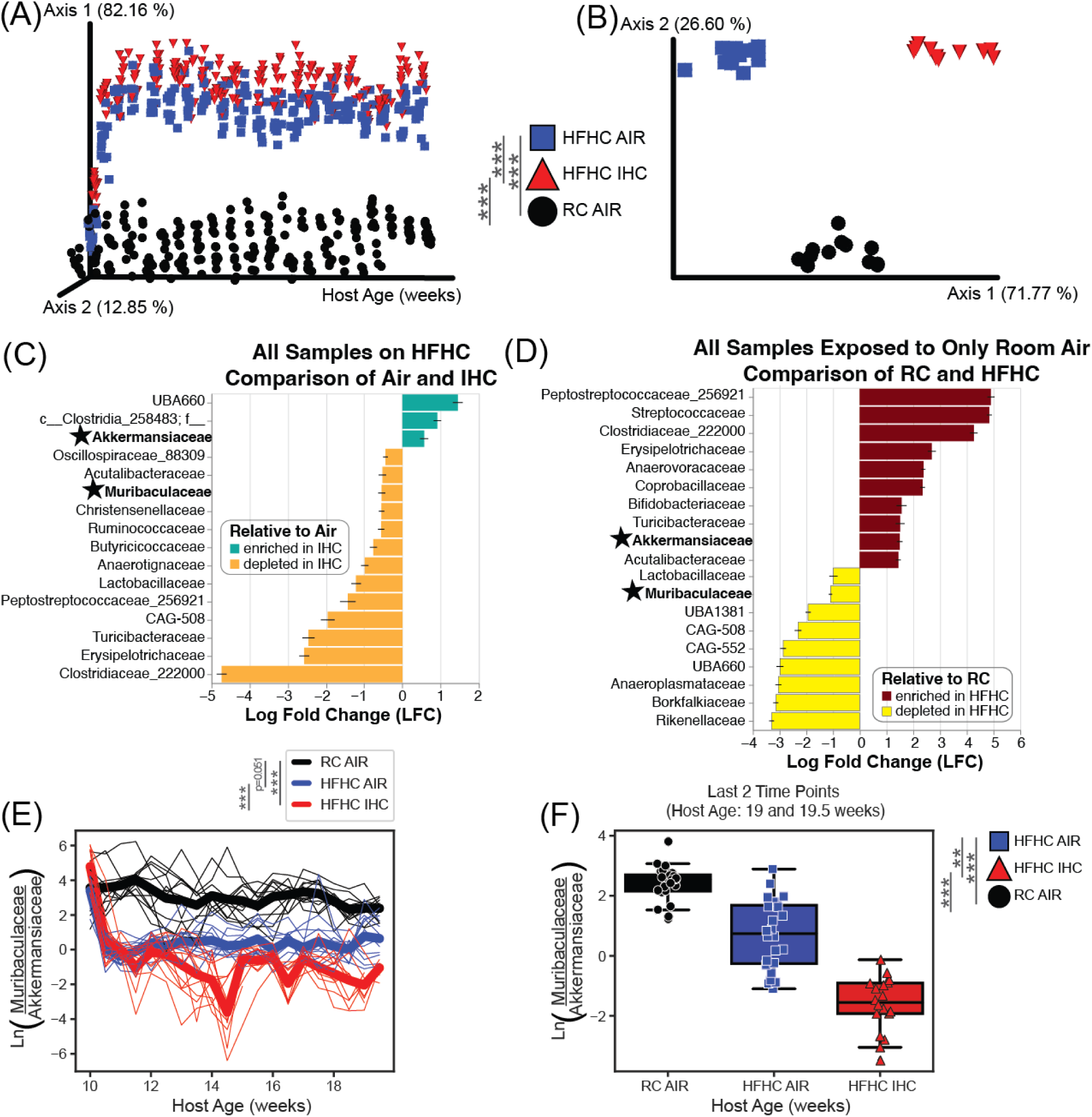
Microbiome analysis of SPF *ApoE*^-/-^ mice on a 10-week HFHC diet with or without IHC treatment. (A) RPCA Beta Diversity PCoA Emperor Plot, metric based on robust center log ratio (rclr) transformation. Each dot represents a single sample. PERMANOVA was used to determine significance. (B) TEMPTED Beta Diversity PCoA of the microbiome. Each point represents a unique mouse. PERMANOVA was used to determine significance. (C) Log fold change barplots the top differentially abundant microbial families in response to exposure type when diet is held constant via ANCOM-BC (p < 0.0001). (D) Log fold change barplots of the top differentially abundant microbial families in response to diet when exposure type is held constant via ANCOM-BC (p < 0.0001). (E) Natural Log Ratio of two families of bacteria previously found to be important in this murine model. The numerator contains all ASV assigned to the Muribaculaceae family; the denominator contains all the ASV assigned to the Akkermansiaceae family. The thick lines represent the mean of all mice in the group and the thin lines represent individual mice over time. Linear mixed effect model (equation: log_ratio ∼ host_age * diet_exp + (1|host_subject_id)) was used to determine significant differences across time. (F) The same log ratio as in C, but only the last 2 time points (host age 19 and 19.5 weeks) shown by group. Mann-Whitney-Wilcoxon test with Bonferroni correction was used to determine significance. Significance: * p < 0.05; ** p< 0.01; *** p < 0.001.

### HFHC and IHC profoundly affect fecal metabolic profiles

Untargeted metabolomics analysis revealed clear separation of the fecal biochemical profiles by both diet and gas exposure via PCA after the exclusion of the first time point, when all animals were on RC diet (Figure 3A). To identify the effect of the HFHC diet on the fecal metabolome, a supervised PLS-DA model was generated comparing the RC-Air and HFHC-Air groups. The model displayed a perfect discrimination (CER = 0) and 3120 features, with VIP score > 1, were considered to be affected by diet (Supplementary Table S3). HFHC diet induced an increase of more than 200 annotated or putatively annotated bile acids, including cholic acid (CA), deoxycholic acid (DCA), lithocholic acid, taurocholic acid, taurodeoxycholic acid, glycocholic acid, 12-ketodeoxycholic acid and their isoforms, as well as the recently described microbial bile acids Ile/Leu-CA, Lys-CA, Phe-CA, Tyr-CA, Val-CA, Phe-DCA, and Glu-DCA. Additionally, HFHC increased the recently described long-chain N-acyl amides, such as Leu-C16:0, Lys-C16:0, Phe-C16:0, Tyr-C16:0, Leu-C18:1, Lys-C18:1, Phe-C18:1, Tyr-C18:1, and Leu-C20:4, and several phosphatidylcholines, including PC(16:0/0:0), PC(16:1/0:0), PC(16:0/20:5), PC(19:0/0:0), PC(17:1/0:0), PC(20:1/0:0), PC(20:3/0:0), PC(15:0/16:0), PC(20:4/0:0), PC(20:2/0:0), PC(14:0/20:4), and PC(15:0/16:0), and sphingosines (both C17 and C18). On the other hand, the HFHC diet appeared to decrease the abundance of more than 50 putative not annotated mono-, di-, tri-, tetra-, and penta-hydroxylated bile acids, several short-chain N-acyl amides, such as cadaverine-C2:0, His-C2:0, acetyl cadaverine-C2:0, His-C3:0, serotonin-C2:0, spermidine-C:20, cadaverine-C4:0, together with indole metabolites, such as 2-oxindole-3-acetic acid, kynurenic acid, tyrosine, and phenylalanine, linoleic acid metabolites, and more than 100 di- and tri-peptides. The log ratio of these extracted features separated animals receiving or not the HFHC diet after the first timepoint, when all animals were receiving RC (Figure 3B). To investigate the effect of the IHC, a supervised PLS-DA model was then generated comparing the HFHC-Air and HFHC-IHC groups. The model displayed a close-to-perfect discrimination (CER = 0.01), with a total of 2034 features affected by gas exposure (Supplementary Table S4). IHC caused the increase of more than 100 annotated and putative bile acids including taurolithocholic acid, taurodeoxycholic acid, taurocholic acid, and 12-ketodeoxycholic acid, several of the above mentioned long-chain N-acyl amides, such as Leu-C16:0, Phe-C16:0, Leu-C18:1, and Phe-C18:1, phosphocholines including PC(16:1/0:0), PC(17:1/0:0), PC(19:1/0:0), PC(20:1/0:0), and bilirubin and urobilin. IHC also reduced the abundance of several bile acids, like cholic acid and microbial bile acids, such as Arg-CA, Glu-CA, Gln-CA, Lys-CA, Glu-DCA, Tyr-DCA, Phe-DCA, short-chain N-acyl amides including serotonin-C2:0 and spermidine-C2:0, vitamine B5, sphingosine C18, and linoleic acid metabolites. The log ratio of RC-Air associated features over HFHC-IHC associated ones significantly separated the three groups of interest (Figure 3C). A comprehensive molecular network of the features affected by HFHC, IHC, or both, was generated (Supplementary Figure 3, interactive Cytoscape file available on GitHub) and sub-networks of molecular features of interest were extracted to showcase some of the molecules driving the separation between the groups, such as bile acids, phosphocholines, and N-acyl amides (Figure 3D). Since atherosclerotic formation was promoted by HFHC and further accelerated by IHC, a subset of 441 features with the concordant or inverse pattern of changes were identified, as these molecules could possibly be biologically highly relevant to the correlated atherosclerosis development (Supplementary Table S5). Representative molecules of interest were also investigated via univariate analysis to highlight group differences (Figure 3E).

**Figure 3.**
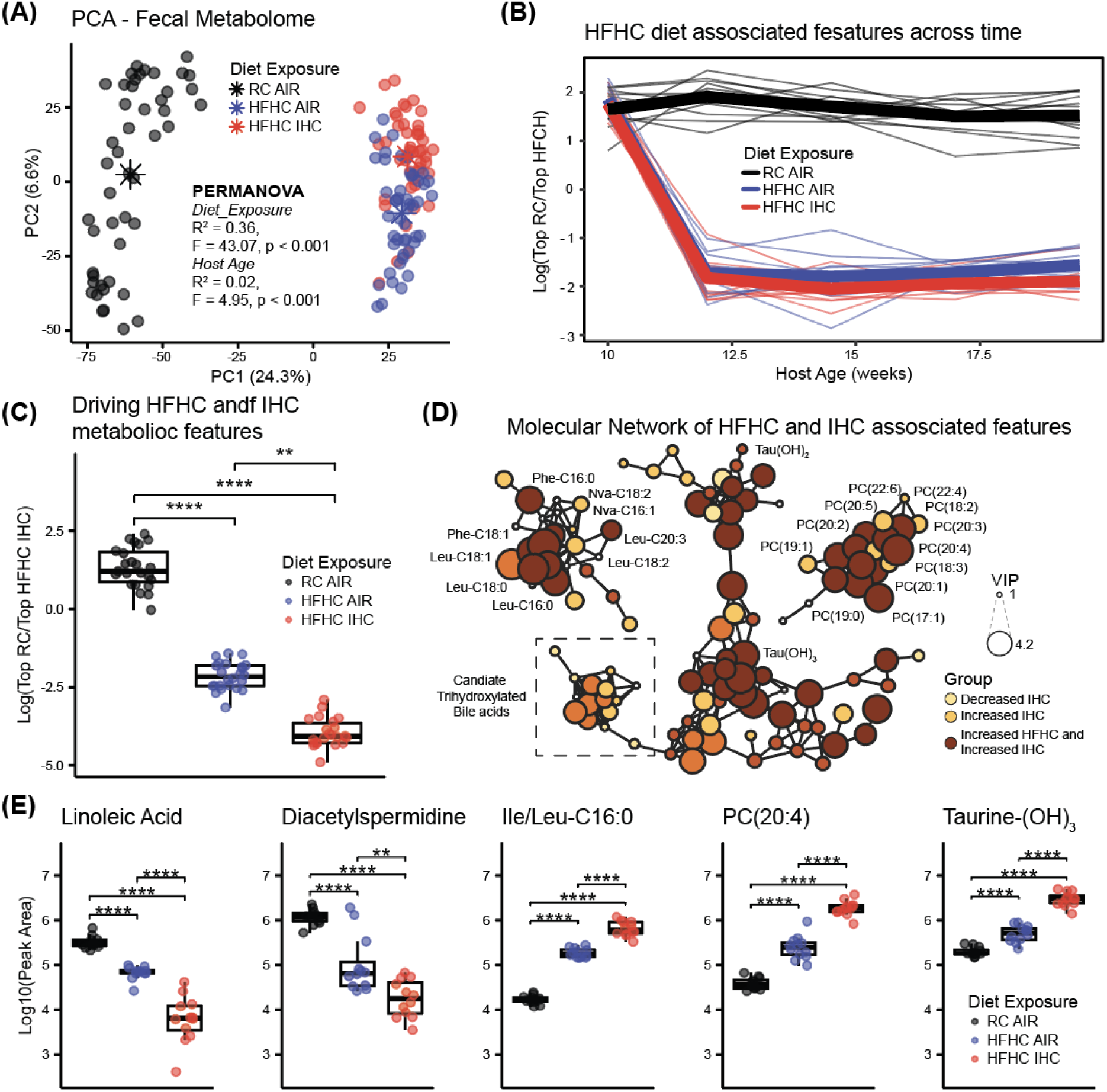
Metabolomics analysis of SPF *ApoE*^-/-^ mice on a 10-week HFHC diet with or without IHC treatment. (A) PCA highlights the separation between groups receiving or not the HFHC diet or being exposed or not to IHC. Centroid separation (indicated with asterisks) was tested using PERMANOVA (Diet_Exposure; R^2^ = 0.36, F = 43.07, p < 0.001). A small effect of host age was observed. (B) Longitudinal analysis of the fecal metabolic profiles using the log ratio of the significant features (3120) associated either with the HFHC (1548) or RC (1572) diet via PLS-DA model (CER = 0). Significance tested using a linear mixed effect model (Log ratio ∼ host_age + diet_exposure + (1|host_subject_id)). (C) Log ratio of selected features (441) significantly associated with either RC-Air (179) or HFHC-IHC (262) via stratified PLS-DA models. Significant separation between the three groups was calculated via ANOVA followed by Tukey’s HSD test. Only the last two timepoints were retained for analysis. (D) Subnetworks of molecular features of interest affected by both HFHC and IHC, i.e. bile acids, phosphocholines, and N-acyl amides. Node sizes are based on VIP scores obtained from the stratified PLS-DA models. (E) Univariate analysis of selected molecular features of interest covering linoleic acid metabolism, polyamines, long-chain N-acyl amides, phosphocholines, and bile acids. Repeated measures were collapsed to mean values across time and the first time point was excluded. Boxplots represent first (lower), interquartile range (IQR), and third (upper) quartile. Whiskers represent 1.5 * IQR. Significance: ** p < 0.01; **** p < 0.0001.

### Akkermansiaceae/Muribaculaceae ratio drives metabolite profiles

The microbiome and metabolome datasets were integrated using joint-RPCA, an unsupervised machine learning model that looks at the joint-factorization and correlation of individual features. Output was filtered to retain only ASVs found to be differentially abundant and used for previous log ratios (Figure 2 and Figure S2). Retained metabolic features were the ones found to be affected by both diet and air exposure via the previously described PLS-DA models (shown in Figure 3C, detailed in Supplementary Table S5). The joint-RPCA co-variance of the top ASV and metabolites of interest show strong patterns (Supplementary Figure S4).

Using the previously discussed microbiome (Figure 2E-F) and metabolome (Figure 3C) log ratios, we used to create a scatterplot to observe sample clustering (Figure 4A). When the log ratios favor the metabolites associated with RC Air conditions and are skewed more towards ASVs from the Muribaculaceae family, there is a majority of RC Air samples. Conversely, when the log ratios favor the metabolites associated with HFHC IHC conditions and are skewed more towards ASVs from the Akkermansiaceae family, there is a majority of HFHC IHC samples. While strong clustering of all three groups is observed, diet appears to have a large effect. In addition, a significant and strong linear relationship can be seen (r=0.80, p=7.6×10^-32^) (Figure 4B). Together, these results indicate that there is a strong microbiome and metabolite signature that appears to be relevant to the response to both diet and exposure type.

**Figure 4.**
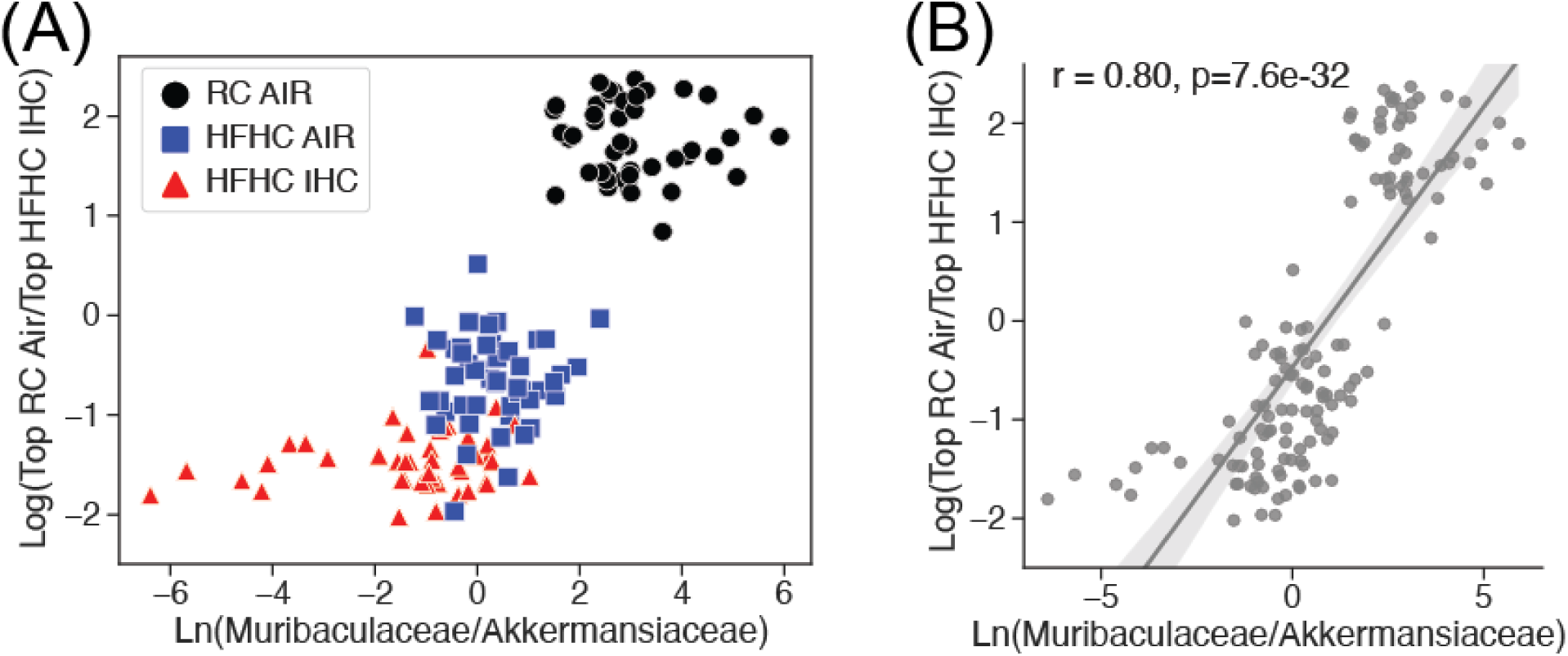
Comparison of microbiome and metabolome log ratios. (A) Scatterplot of microbiome and metabolome log ratios of interest (as described in Fig 2E-F, 3C), colored by experimental group (B) Linear regression plot of data shown in A.

### Absence of microbiota reduces HFHC- and IHC-induced atherosclerosis in the aorta but not in the pulmonary artery

To investigate if gut microbiota plays a role in HFHC- and IHC-induced atherosclerosis, germ-free (GF) *ApoE*^-/-^ mice were fed the same HFHC diet in the presence of either Air or IHC for 10 weeks, as for the SPF animals. The atherosclerotic lesions of GF mice were compared to corresponding sex- and age-matched conventionally-reared SPF controls. HFHC-induced lesions were significantly lower in the aorta, mainly in the aortic trunk which excludes the arch, in GF mice than in SPF mice (Figure 5A-B, Aorta, GF-HFHC 4.4±0.59% vs SPF-HFHC 6.8±0.47%, p < 0.05; Aortic trunk, GF-HFHC 1.9±0.0.66% vs SPF-HFHC 5.0±0.65%, p < 0.05). Although the aortic arch and pulmonary artery showed a higher lesion formation (8-12%), the differences between SPF and GF were not significant. When exposed to IHC, GF *ApoE*^-/-^ mice showed statistically less lesion only in the aortic trunk relative to SPF controls but not in the other examined vascular areas (Figure 5C-D; Aortic trunk, GF-IHC 1.4±0.45% vs SPF-IHC 5.9±1.49%, p<0.05). These data suggest that microbial colonization modulates atherosclerosis differently in distinct vascular beds.

**Figure 5.**
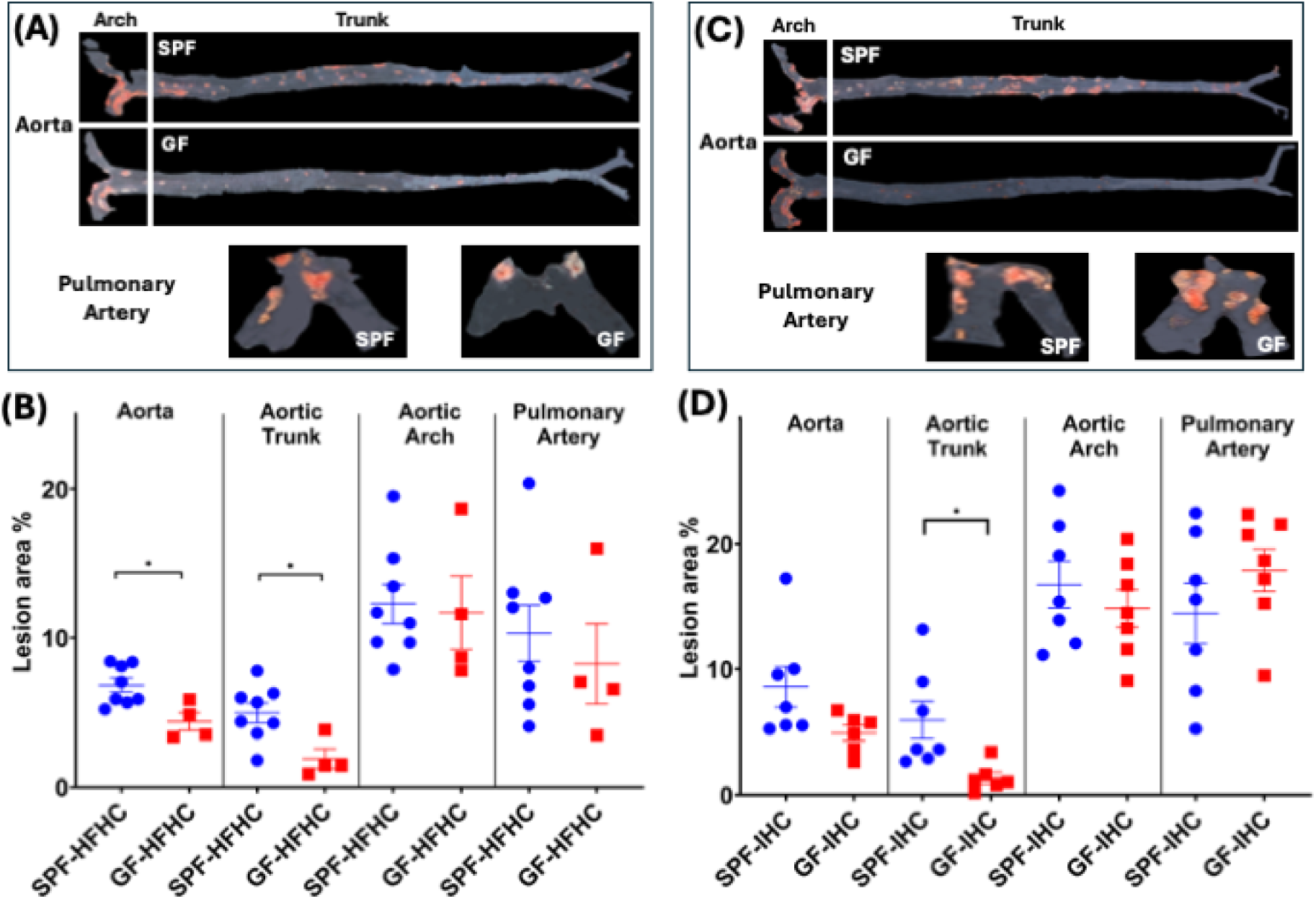
Atherosclerotic lesions in GF and SPF *ApoE*^-/-^ mice, after 10-week of HFHC diet or IHC exposure. (A, C) Representative Sudan IV-stained images of lesions. (B, D) The en-face lesions were quantified as the percentage of lesion area in the total area of the blood vessel examined. Atherosclerotic lesions were significantly reduced in the aorta and aortic trunk of HFHC-fed GF mice and aortic trunk of IHC-exposed GF mice. Data were presented as means ± SEM. Significance with Student’s t-test, * p<0.05.

As shown in Figure 1, IHC promoted the formation of atherosclerosis in the aorta, aortic arch, and pulmonary artery of SPF *ApoE*^-/-^ mice in the presence of HFHC. However, the effect of IHC on atherosclerosis was only detected in the pulmonary artery not in the aorta of GF *ApoE*^-/-^ mice fed with HFHC (Figure 6; PA, GF-IHC 17.9±1.69% vs GF-Air 8.3±2.70%, p = 0.01). These data indicate that (1) microbiota is required for IHC-induced atherosclerosis in the aorta; (2) microbiota-independent mechanisms underlie IHC-induced atherosclerosis in the pulmonary artery.

**Figure 6.**
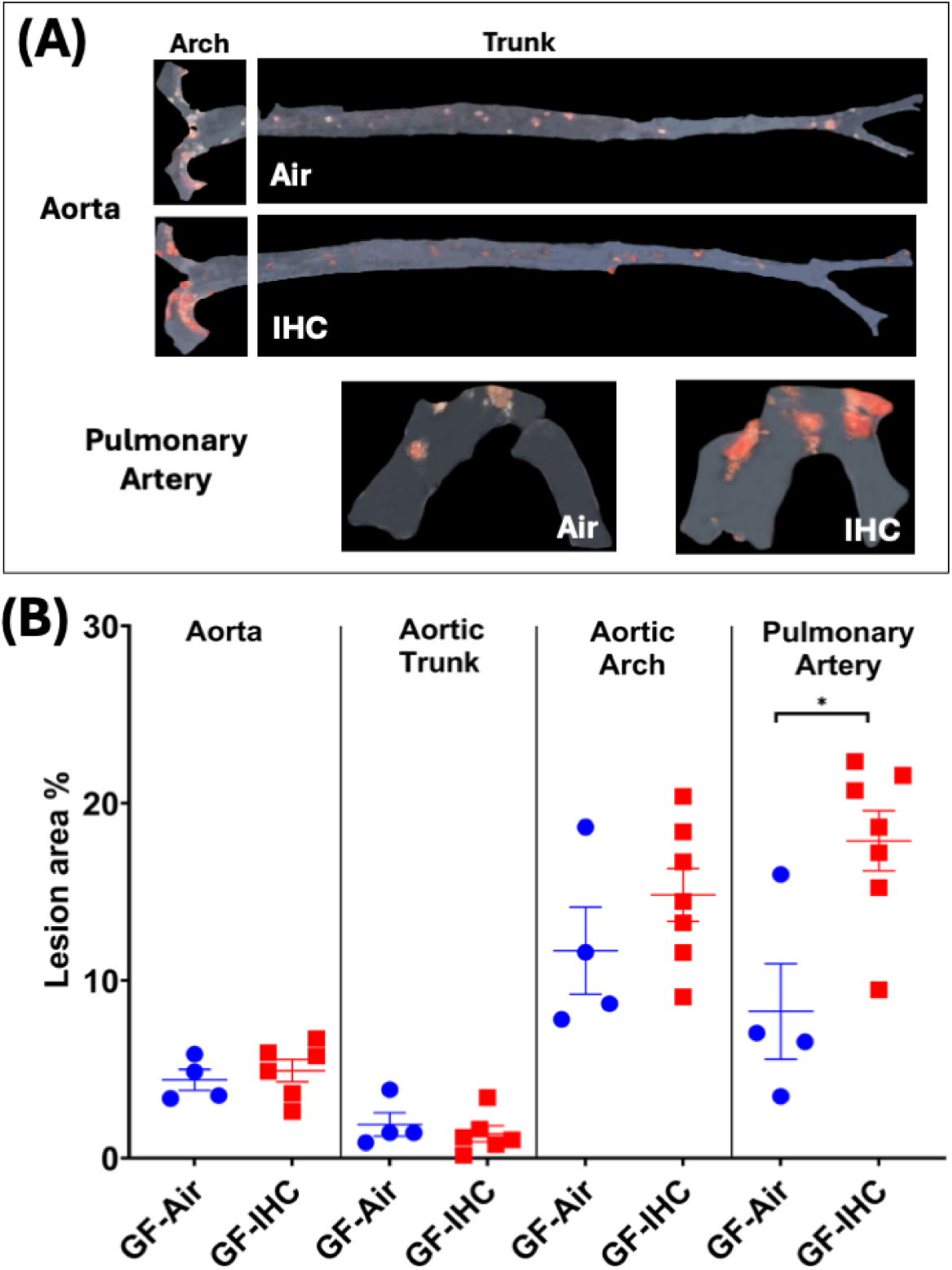
Atherosclerotic lesions in GF *ApoE*^-/-^ mice after 10-week IHC and Air treatment. (A) Representative Sudan IV-stained images of lesions. (B) The en-face lesions were quantified as the percentage of lesion area in the total area of the blood vessel examined. In the absence of microbiota, IHC exposure promoted the progression of atherosclerosis in the pulmonary artery but not in the aorta. Data were presented as means ± SEM. Statistical significance with Student’s t-test, * p<0.05.

## Discussion

OSA is a sleep disorder characterized by repetitive episodes of complete or partial airway obstruction, resulting in IHC which is a prominent feature of OSA pathophysiology. The most commonly used experimental IH protocol is implemented by rapid delivery of a hypoxic mixture to an airtight chamber followed by flushing with normoxic air. Reducing the ambient chamber oxygen to 5–10% results in an SaO_2_ of 60–80% [53]. IH exposure is given during daytime when rodents generally sleep. Since OSA can vary in severity from few (i.e., 5) to many (i.e., 15-30) episodes per hour, we chose a middle of the range treatment that is practically achievable in the present study. We also included hypercapnia in our experimental protocol to better mimic the oscillations of O_2_ and CO_2_ in OSA patients.

In the current study, we investigated the impact of a high fat and high cholesterol diet, as well as its interaction with IHC, on the fecal microbiome and metabolome in relation to the development of atherosclerosis using both conventionally-reared SPF and GF *ApoE^-/-^* mice. High fat and high cholesterol are two common components in atherogenic diets to induce and facilitate atherosclerosis in experimental mice [54, 55]. *ApoE^-/-^* mice are a widely used mouse model for atherosclerosis research because *ApoE*-deficiency leads to increased plasma levels of total cholesterol (mostly in the chylomicron remnant/VLDL fractions) and increases sensitivity to dietary lipids [56]. In the present study, we demonstrated that the HFHC diet significantly promoted atherosclerosis in the aorta and pulmonary artery of *ApoE*^-/-^ SPF mice (Figure 1). We also performed another study using a high fat only diet (HF) and found that the HF diet alone caused much less lesions (<5%), which was similar to the RC control group. Moreover, IHC, in the absence of high cholesterol, increased atherosclerosis but the extent of lesion was still minimal. (Supplementary Figure S5). This set of results demonstrated that high cholesterol, rather than high fat, is essential for atherogenesis in the *ApoE^-/-^* mice under both Air and IHC conditions. When exposed to IHC, atherosclerotic progression was expedited in addition to the HFHC effect (Figure 1). It is worth mentioning that IHC stress reduced body weight gain and food intake (Supplementary Figure S1). Even though consuming less HFHC food, IHC mice still developed more lesions, indicating that IHC exposure itself has atherogenic effect.

The gut microbiome is a complex ecosystem of microorganisms that live in the host intestines. Diet, environmental exposures, stress and disease-relevant stimuli could cause imbalance of these microorganisms (i.e., gut dysbiosis) and subsequently have various impacts on host health. Current microbiome data revealed that the Akkermansiaceae family was enriched and the Muribaculaceae family was depleted under HFHC and IHC conditions (Figure 2C-D). Natural log ratio of these two families separates the three experimental groups well (Figure 2E-F), suggesting that these two families contribute to OSA-induced atherosclerosis. Some species within the Muribaculaceae are thought to help support gut barrier function [57, 58], produce short-chain fatty acids by metabolizing dietary fiber and mucin glycans [59] and its role in mouse intestinal inflammation is controversial [60]. While Akkermansiaceae members, especially *Akkermansia muciniphila*, are known for their mucin degradation enzymes [61, 62], as well as their impact on host lipid and cholesterol homeostasis [63, 64].

Bacterial families Peptostreptococcaceae, Streptococcaceae and Clostridiaceae were relatively increased by HFHC diet (Figure 2D), which corroborates the finding that these bacteria levels were significantly higher in western diet-fed mice with atherosclerosis [65]. Clostridiaceae and Peptostreptococcaceae are more abundant in omnivores and associated with higher trimethylamine N-oxide (TMAO) levels in humans [66], suggesting their potential role in metabolism of dietary choline, phosphatidylcholine, and L-carnitine into trimethylamine (TMA) which is further oxidized to TMAO in the liver. TMAO is a well-known risk factor for atherogenesis by enhancing macrophage cholesterol accumulation and foam cell formation, altering bile acid composition and pool size as well as causing endothelial inflammatory injury [66, 67]. Genus *Streptococcus* belongs to the family Streptococcaceae, its members have been found in human atherosclerotic plaques [68] and its abundance in the gut was associated with coronary atherosclerosis and systemic inflammation [69].

We also identified bacterial families affected by IHC exposure, such as Ruminococcaceae and Lachnospiraceae (Figure 2C, Supplementary Figure S2E and Supplementary Table S1). In terms of atherosclerosis, Ruminococcaceae is reduced in atherosclerosis patients compared to healthy controls [70, 71], negatively associated with cardiometabolic diseases through isolithocholic acid, muricholic acid and nor cholic acid [72] as well as regulates lipid metabolism, apolipoprotein and cholesterol [73]. Lachnospiraceae is positively correlated with total and LDL cholesterol and its abundance decreases in patients with coronary artery disease (CAD) [74]. Kasahara et al. discovered that *Roseburia intestinalis*, a member of the Lachnospiraceae family, interacts with dietary plant polysaccharides to produce butyrate and protect against atherosclerosis by improving intestinal barrier function thus lowering systemic inflammation and directing metabolism away from glycolysis and toward fatty acid utilization [75]. Our current microbiome data have actually demonstrated that both Ruminococcaceae and *Roseburia intestinalis* from the Lachnospiraceae family were depleted by IHC (Figure 2C, Supplementary Figure S2E and Supplementary Table S1), implicating their roles in mediating IHC-induced atherosclerosis.

Interestingly, bile acids (BAs) appeared to be top differentially abundant metabolites under HFHC and IHC conditions (Supplementary Tables S3 and S4). Bile acids are synthesized from cholesterol in the liver and prevent cholesterol overload in the body, therefore ameliorate atherosclerosis. BAs also act as signaling molecules and work through bile acid receptors (BARs), such as nuclear receptors—farnesoid X receptor (FXR) and membrane-bound receptors—Takeda G protein-coupled receptor (TGR5), to regulate their own homeostasis, lipid, glucose and energy metabolism, gut barrier integrity, inflammation as well as cardiovascular function [76]. It has been reported that FXR and TGR5 agonists have lipid-lowering and anti-inflammatory effects [76]. Hence, the changed bile acid by HFHC and IHC can serve as either agonists or antagonists of FXR or/and TGR5 receptors to modulate atherosclerotic formation.

In addition, HFHC- and IHC-induced gut dysbiosis can affect bile acid composition/pool size and indirectly influence host functions via BA signaling. Three major microbial enzymes, i.e. bile salt hydrolases (BSHs), hydroxysteroid dehydrogenases (HSDHs), and bile-acid-inducible (bai) genes, are responsible for the generation of various secondary BAs. Notably, among the altered bacterial families by HFHC and IHC (Figure 2C-D, Supplementary Figure S2E-F, Supplementary Tables S1 and S2), Ruminococcaceae, Lachnospiraceae and Peptostreptococcaceae contain bai genes which encode enzymes involved in 7α-dehydroxylation [77]. Lactobacillaceae, Streptococcaceae and Clostridiaceae carry BSH genes which encode enzymes catalyzing the deconjugation of the N-acyl amide bond between primary BAs and taurine or glycine [76]. Our integrated analysis of the microbiome and metabolome uncovered that different metabolite signatures were correlated with Muribaculaceae and Akkermansiaceae families (Figure 4 and Supplementary Figure S4). It is unclear at present if the microbial families are actively involved in altering the amounts or/and activities of these metabolites and this requires further investigation.

Our previous and present studies have demonstrated that IHC exposure significantly alters both gut microbiota and bile acids along with an elevated level of atherosclerosis in the HFHC fed mice [25, 26, 52, 78]. Hence, it is critical to examine the synergistic role of the gut microbiota, HFHC diet, and IHC on atherogenesis. Our current data revealed that in the absence of microbial colonization, HFHC diet-stimulated development of atherosclerosis was significantly attenuated in the aorta, mostly in the aortic trunk, under both room air and IHC conditions (Figure 5), suggesting that the microbiome contributes to atherosclerotic disease pathogenesis. Although the underlying mechanisms remain elusive, the gut microbiota might mediate HFHC diet- and IHC- induced atherogenesis through: (1) activating host’s immune system (such as activation of macrophages), (2) elevating the inflammatory response (such as releasing lysophospholipids (LPS) and peptidoglycan (PG)), (3) catalyzing cholesterol metabolites (such as the production of secondary bile acids).

Unlike the aorta, the IHC-induced atherosclerosis was not abolished in the absence of microbial colonization in the pulmonary artery (PA) (Figure 6), suggesting that the gut microbiota is not a major player. Other mechanisms may be responsible for pathogenesis of PA atherosclerosis when exposed to IHC, such as an increase of shear stress at branching points of the pulmonary arteries [79] and/or pulmonary hypertension induced by intermittent hypoxia [52], with both potentially causing endothelial damage and initiating atherosclerotic formation. Future research is warranted.

## Conclusion

Taken together, we demonstrated that IHC and HFHC work synergistically to promote atherosclerosis, which requires the presence of gut microbiota. Bacterial families Akkermansiaceae and the Muribaculaceae, as well as microbial metabolite bile acids appear to be important modulators of OSA-induced atherosclerosis. The knowledge obtained helps us better understand the mechanistic link between diet, microbiota and IHC/OSA-induced atherosclerosis and provides a basis for future therapeutic approaches to prevent and treat OSA-induced atherosclerosis, such as prebiotics, probiotics or synbiotics to correct gut dysbiosis or FXR/TGR5 agonists or antagonist to modify BA signaling, combined with diet modification.

## List of Abbreviations

ApoE: apolipoprotein E
HCHF: high cholesterol high fat
RC: regular chow
PA: pulmonary artery
GF: germ-free
SPF: conventionally reared specific pathogen free

## Data Availability

Microbiome data is available under the EBI accession number ERP110592. Code for microbiome and joint-RPCA analysis can be found on GitHub (https://github.com/knightlab-analyses/OSAstudy_16S_HFHC). Metabolomics data is available in GNPS/MassIVE under the accession code MSV000082972 (https://massive.ucsd.edu/ProteoSAFe/dataset.jsp?task=7996239533ea48738b550650e8531142) and code to analyze data and generate figures present in the manuscript is available on GitHub (https://github.com/simonezuffa/Manuscript_HFHC_IHC).

## Author Contributions

J.X. and G.G.H. conceived and designed research; O.P. and J.M. performed experiments; J.X., C.A., S.Z., O.P. and D.Z. analyzed data; J.X., C.A., S.Z., D.Z. and G.G.H. interpreted results of experiments; J.X., C.A., S.Z., O.P. and D.Z. prepared figures; J.X., C.A. and S.Z. drafted manuscript; J.X., D.Z., P.D., R.K. and G.G.H. edited and revised manuscript; J.X., C.A., S.Z., O.P., J.M., D.Z. P.D., R.K. and G.G.H. approved final version of manuscript.

## Acknowledgements

We thank Dr. Sarkis Mazmanian’s laboratory at CalTech for rederivation of the germ-free *ApoE*^-/-^ mice.

## Declarations

### Ethics approval and consent to participate

All animal protocols were approved by the Animal Care Committee of the University of California San Diego and followed the Guide for the Care and Use of Laboratory Animals of the National Institutes of Health.

### Consent for publication

Not applicable.

