## Supplementary Figures and Legends for "Gut Microbiota and Derived Metabolites Mediate Obstructive Sleep Apnea Induced Atherosclerosis"

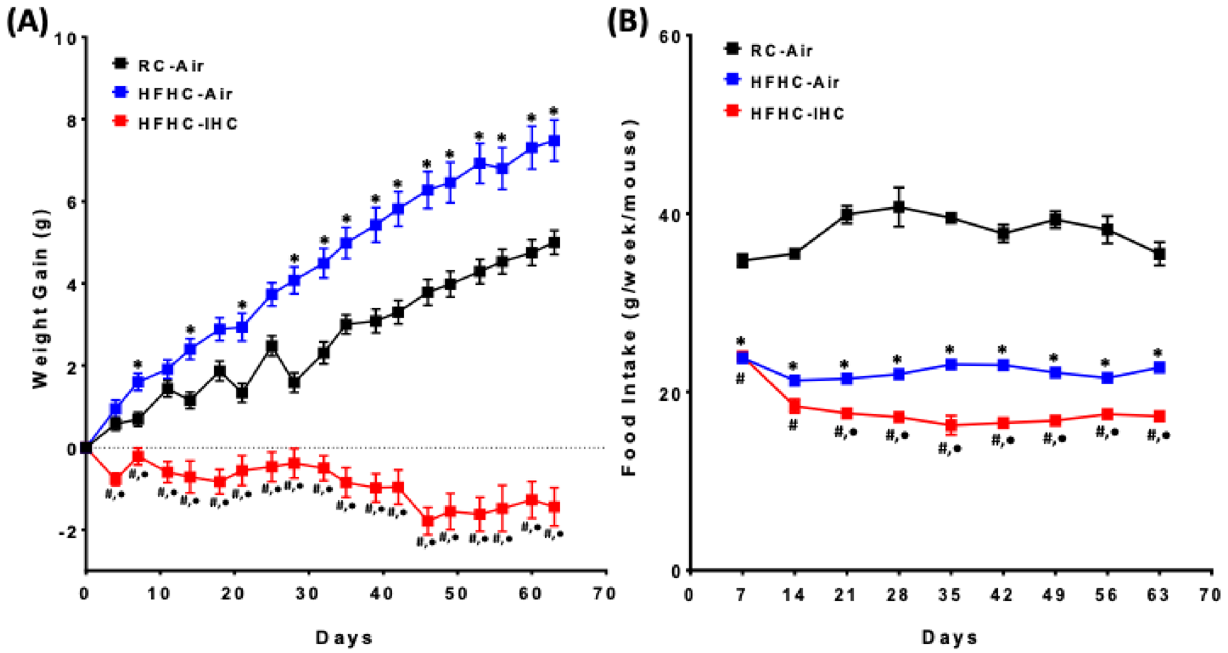


**Supplementary Figure S1. (A) Body weight gain and (B) Food consumption.** RC-Air vs HFHC-Air vs HFHC-IHC. Mice on the HFHC diet showed greater weight gain than those in the RC group. IHC with HFHC caused weight loss over the treatment time. More RC food was consumed than HFHC food. Under the same HFHC condition, food intake was reduced by IHC exposure starting week 3 of the treatment. Multiple t-tests with Bonferroni correction, $P<0.05$, * HFHC-Air vs RC-Air, # HFHC-IHC vs RC-Air and · HFHC-IHC vs HFHC-Air.

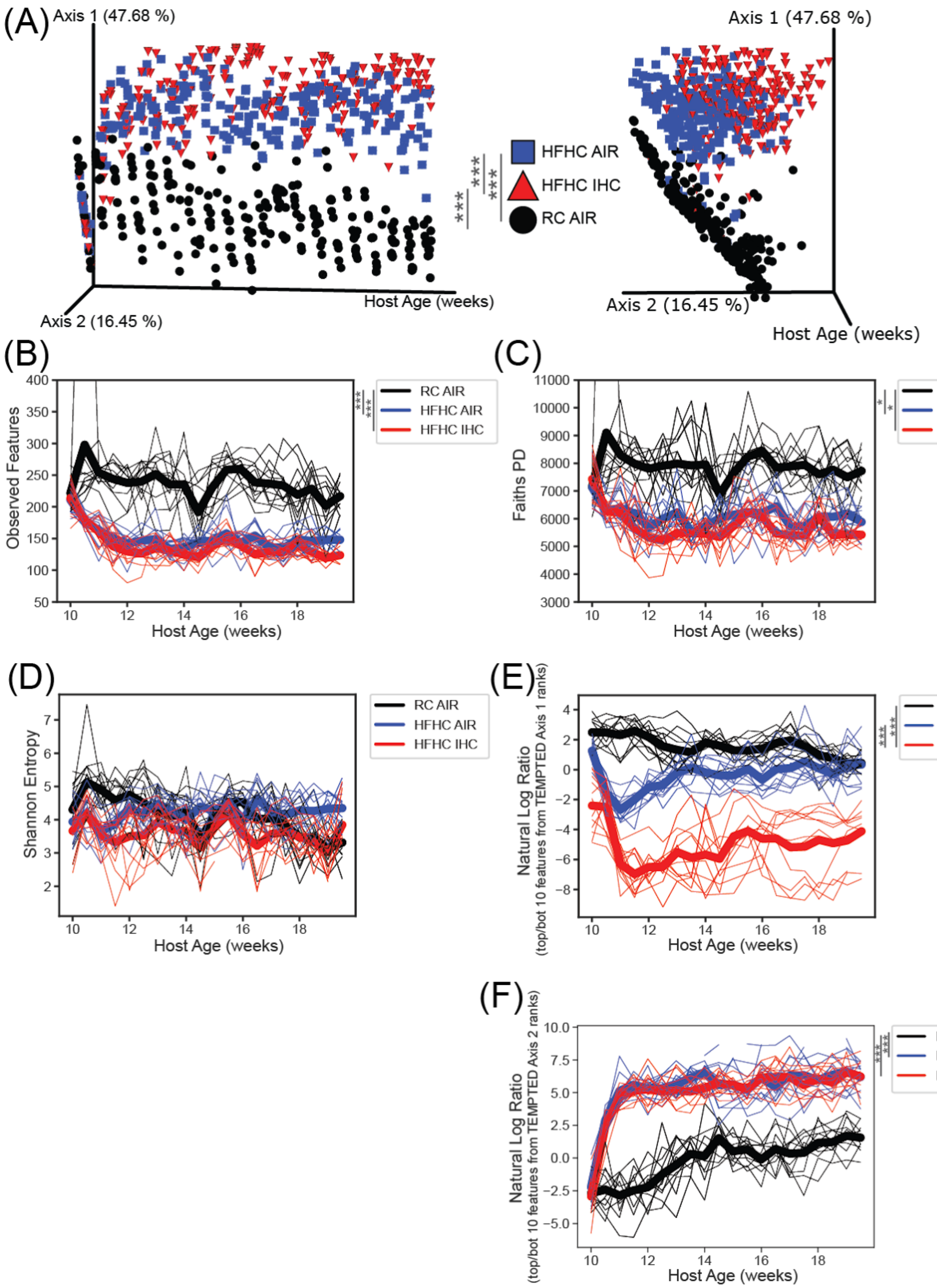


**Supplementary Figure S2. Additional 16S V4 microbiome analysis.** (A) Weighted UniFrac Beta Diversity PCoA metric takes into account abundance and phylogeny. Each dot represents a single sample. Lateral view on left, end-on view on right. PERMANOVA was used to determine significance. Alpha Diversity Metrics: (B) Observed Features (unique ASVs) (C) Faith's Phylogenetic Diversity, and (D) Shannon Entropy. The thick line represents the mean of all mice in the group and the thin lines represent individual mice over time. Linear mixed effect model (equation: alpha diversity metric value ~ host_age * diet_exp + (1|host_subject_id)) was used to determine significant differences across time. (E) Natural Log Ratio of the top and bottom 10 differentially ranked ASVs as present in TEMPTED Axis 1. (F) Natural Log Ratio of the top and bottom 10 differentially ranked ASVs as present in TEMPTED Axis 2. See Supplementary Tables S1 and S2 for ASVs and taxonomic annotation. The thick line represents the mean of all mice in the group and the thin lines represent individual mice over time. Linear mixed effect model (log_ratio ~ host_age * diet_exp + (1|host_subject_id)) was used to determine significant differences across time. Significance: ns $p > 0.05$; * $p < 0.05$; ** $p < 0.01$; *** $p < 0.001$.

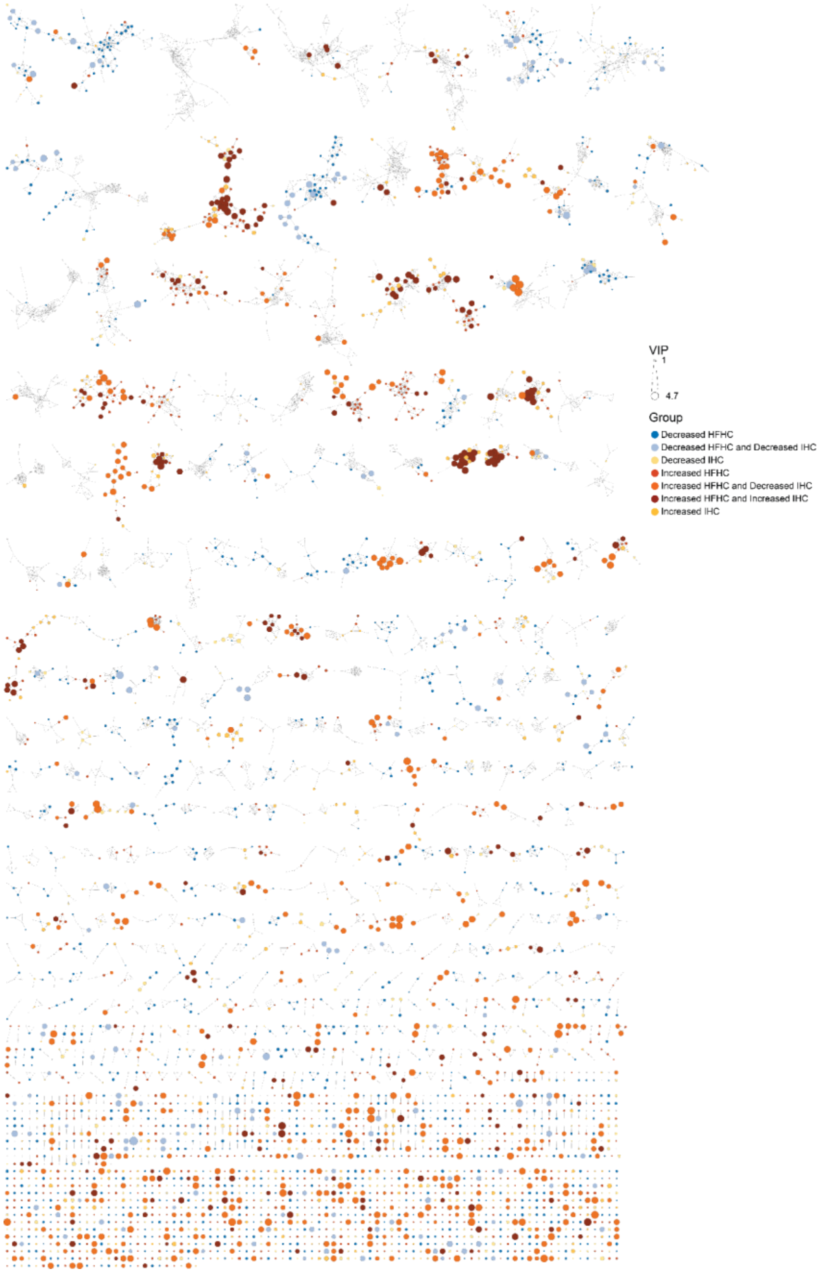


**Supplementary Figure S3**. **Molecular network.** Molecular network filtered for significant features from the generated PLS-DA models. Variable importance represented by size of nodes (VIPs [1,4.7]). Colors indicate groups in which the features were present in higher abundances. A full explorable and interactive Cytoscape file is available to download on GitHub (https://github.com/simonezuffa/Manuscript_HFHC_IHC).

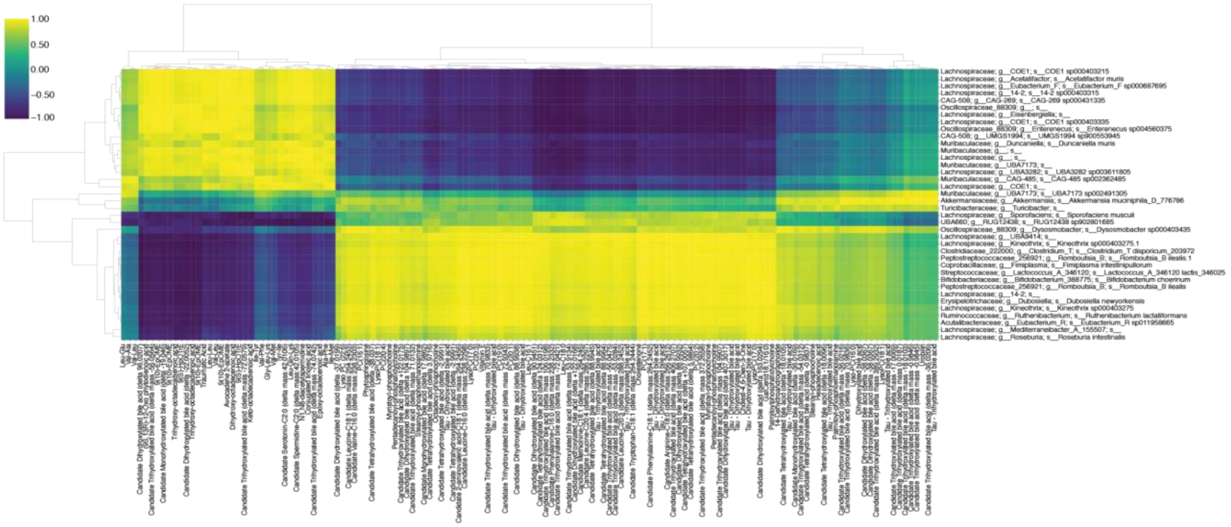


**Supplementary Figure S4. Joint-RPCA of top microbes and metabolites of interest.** All ASVs of interest (y-axis) and used in any of the log ratios (x-axis; Figure 2E-F, S2E-F) and their co-variance with all metabolites of interest (Figure 3C, Supplemental Table 1-2) based on the data from the final time point in common (host age = 19.5 weeks) using joint-RPCA.

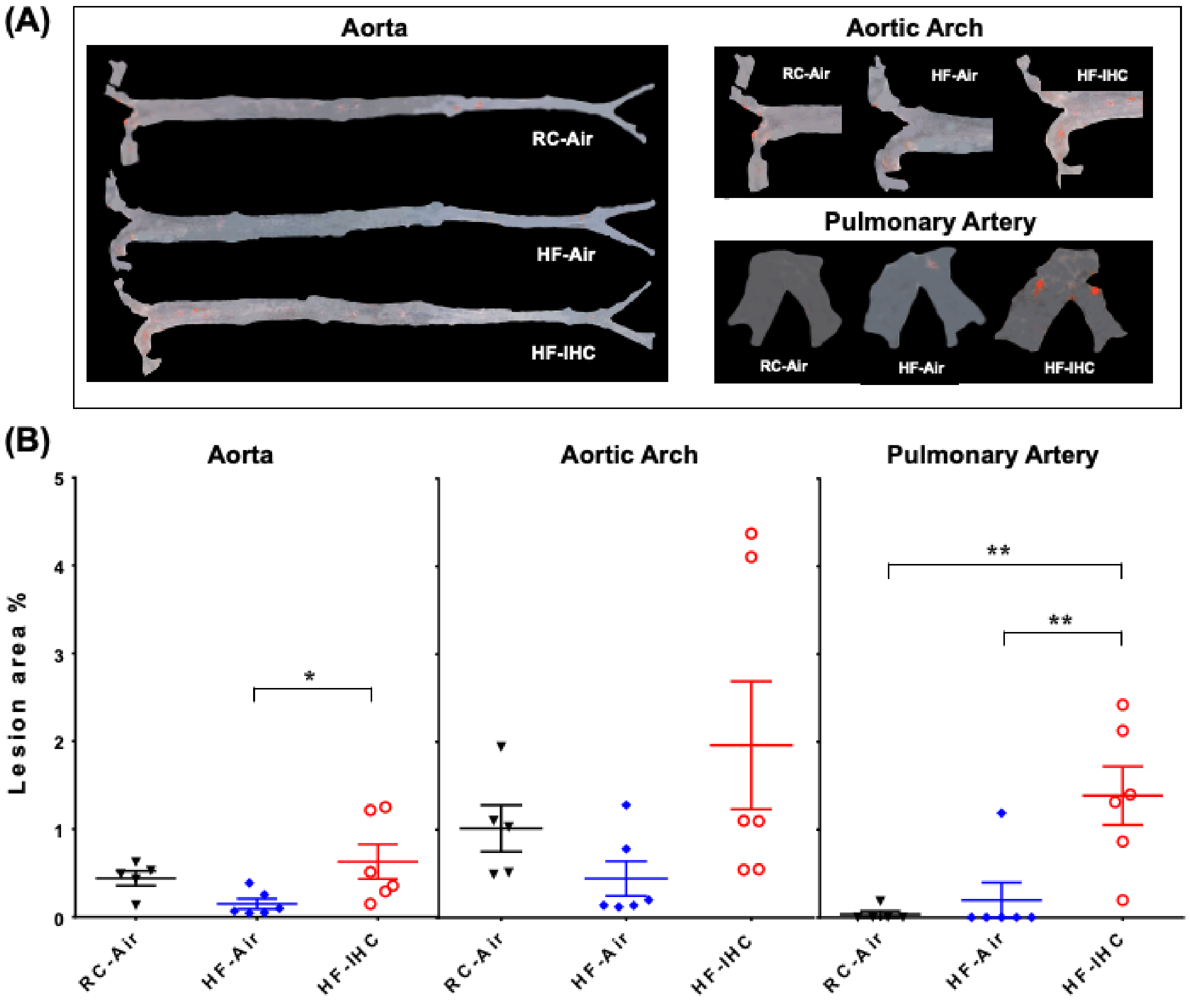


**Supplementary Figure S5. Atherosclerotic lesions after 10-week of high fat (HF) diet with or without IHC in SPF *ApoE*$^{-/-}$ mice.** (A) Representative Sudan IV-stained images of lesions. (B) The en-face lesions were quantified as the percentage of lesion area in the total area of the blood vessel examined. The HF diet caused less than 5% lesion formation. IHC promoted atherosclerotic progression in the presence of HF compared to the controls in the aorta and pulmonary artery. Data are presented as means ± SEM. Statistical significance tested via One-way ANOVA followed by Tukey's multiple comparison test. Significance: *p<0.05, ** p < 0.01.
